# dACC Response to Presentation of Negative Feedback Identifies Stimulant Dependence Diagnoses and Stimulant Use Severity

**DOI:** 10.1101/317990

**Authors:** Eric D. Claus, Matthew S. Shane

## Abstract

Error-monitoring abnormalities in stimulant-dependent individuals (SDIs) may be due to reduced awareness of committed errors, or to reduced sensitivity upon such awareness. The distinction between these alternatives remains largely undifferentiated, but may have substantial clinical relevance. We sought to better characterize the nature, and clinical relevance, of SDIs’ error-monitoring processes by comparing carefully isolated neural responses during the presentation of negative feedback to a) stimulant dependence status and b) lifetime stimulant use. Forty-eight SDIs and twenty-three non-SDIs performed an fMRI-based time-estimation task specifically designed to isolate neural responses associated with the presentation (versus expectation) of contingent negative feedback. SDIs showed reduced dACC response compared to non-SDIs following the presentation of negative feedback, but only when error expectancies were controlled. Moreover, lifetime stimulant use correlated negatively with magnitude of dACC attenuation. While this findings was minimized after controlling for age, these results suggest that SDIs may be characterized by a core reduction in neural activity following error feedback, in the context of intact feedback expectancies. Correlations with lifetime stimulant use suggest that this neural attenuation may hold clinical significance.

## Introduction

Stimulant dependent individuals (SDIs) are characterized by a persistent use of drugs, despite repeated negative consequences, and this has been interpreted in terms of a potentially deficient action-monitoring system (Franken et al., 2007; Hester et al., 2013; 2007). A growing body of work has provided both electrophysiological and hemodynamic evidence indicating that substance abuse is characterized by abnormal ACC/mFC responses to rewards and errors (Hester et al., 2013; Kaufman et al., 2003; Steele et al., 2014); but see (Alexander et al., 2015; Baker et al., 2016; Castelluccio et al., 2013), as well as to feedback indicating rewards and errors (Fedota et al., 2015; Parvaz et al., 2015; Patel et al., 2013). Moreover, a handful of recent studies have demonstrated significant relationships between magnitude of dACC/mFC response to error-related information and clinically-relevant downstream metrics of abuse (Luo et al., 2013; Marhe et al., 2013; Moeller et al., 2014; Steele et al., 2014). Nonetheless, the precise nature of these error-monitoring abnormalities is still unclear. For instance, it remains to be determined whether SDIs are characterized by a reduced identification of committed errors, or to a reduced sensitivity upon such identification. Work in normative populations has consistently validated this distinction (e.g. (Delgado et al., 2000), and has highlighted its importance for characterizing the nature of error-monitoring failures when they occur. Applied to SDIs, evidence of reduced identification of errors may imply a central impairment in the ability to recognize relevant behavioral outcomes (Hester et al., 2007). Alternately, evidence of an attenuated neural response subsequent to error-identification may imply a more basic insensitivity to the presentation of error-related information. These subtle differences in the nature of SDI’s error-monitoring abnormalities may thus underlie important details regarding the pathogenesis of the disorder, and may help inform the development of targeted treatment protocols.

Of significant inconvenience, precise isolation of neural responses underlying the presentation of error-related feedback is not so easily accomplished, as the magnitude of neural response to error are generally reciprocal to that error’s prior expectancy (Holroyd and Coles, 2002; Holroyd et al., 2004); thus, the mere expectation of error-related feedback may interfere with the accurate characterization of presentation-phase sensitivity. Several approaches devised to reduce the impact of this reciprocity have attempted to remove the contingency between response and outcome (e.g. guessing tasks, random-outcome gambling tasks; (Delgado et al., 2000); however, in so doing, these tasks have difficulty evaluating critical relationships between outcome processing and subsequent behavioral change (Shane and Weywadt, 2014). Other approaches have parsed the ‘expectation’ versus ‘presentation’ phases of outcome processing into distinct temporal units (e.g. monetary incentive delay tasks; MID; (Knutson et al., 2001a; 2001b; 2005)); while this retains important response-outcome contingencies, it is unclear whether the temporal isolation of expectation/presentation phases truly controls for the reciprocal nature of error-related activity during each phase (Shane and Weywadt, 2014).

Perhaps due to these shortcomings, only three studies have attempted isolation of presentation-phase activity in SDIs and these have yielded inconsistent results. Patel et al. (2013) failed to find cocaine-related dACC reductions during either phase of a MID task. Similarly, Rose et al. (2017) failed to find any differences in dACC in cocaine users compared to controls, but did report reduced reward outcome related responses in the right habenula in cocaine users. In contrast, a more recent study reported that cocaine-dependent individuals showed electrophysiological evidence of reduced sensitivity to unexpected, but not expected, error feedback on a gambling-type task (Parvaz et al., 2015). The latter result suggests that SDIs’ error-monitoring abnormalities may be caused by fundamental insensitivity to the presentation of error-related feedback. However, additional research would aid firm conclusions, and help determine the extent to which reduction in error-sensitivity are associated with downstream metrics of abuse.

To these ends, the present study utilized an fMRI-based time-estimation task specifically designed to evaluate neural responses following the presentation of outcome feedback within an environment that controlled for prior expectancies (Shane and Weywadt, 2014). Of primary interest was the extent to which SDIs and non-SDIs would show differential neural responses following presentation of error feedback when variance associated with prior expectations was experimentally controlled. Consistent with the extant literature (Hester et al., 2013; Kaufman et al., 2003; Parvaz et al., 2015), we hypothesized that SDIs would demonstrate reduced error-related responses compared to non-SDIs following presentation of contingent negative feedback. Of import, and consistent with Parvaz et al (2015), we predicted that this reduced error-related response would occur only when error-expectancies were controlled, but would disappear when error-expectancies were allowed to vary. This result would suggest that SDIs’ error-monitoring abnormalities are the result of reduced sensitivity to unexpected negative feedback, rather than by reduced tendency to form such expectations.

An additional goal of the present study was to serve as a preliminary test of clinical utility of any witnessed error-related abnormalities in the SDI group. Only a handful of such studies have been conducted to date, and none has attempted to parse expectation/presentation phases of outcome processing (Parvaz et al., 2015; Patel et al., 2013). With this in mind, we evaluated the extent to which the magnitude of error-related responses would relate to lifetime stimulant use (Parvaz et al., 2015). Following our broader hypothesis, we predicted that these clinically-relevant relationships would also present only when error-expectancies were controlled.

## Methods and Materials

### Participants and Diagnostic Categories

Eighty-four participants (were recruited through advertisements placed in probation/parole offices throughout the Albuquerque metropolitan area. Advertisements specifically targeted individuals with a criminal history who were currently on probation or parole with or without a history of drug use. Participants were phone-screened regarding key inclusion/exclusion components and further screened in person to determine eligibility. After obtaining consent, SDI status was determined via the Structured Clinical Interview for DSM-IV Disorders (SCID; (First et al., 2002)). SDI met dependence criteria for cocaine or stimulant dependence; non-SDI met criteria for neither. To best compare SDI and non-SDI groups, and isolate differences specifically associated to stimulant dependence, both SDI and non-SDI groups had other co-morbid abuse/dependence diagnoses. Thus, any differences seen between groups can be more confidently attributed to the stimulant dependence diagnosis rather than some other comorbid condition. Exclusionary criteria included: history of head injury that caused >30 minutes lost consciousness, past or current history of CNS disease (e.g. stroke, MS, epilepsy or repeated seizure history of any kind, tumor, etc.) or brain lesion, Axis I DSM-IV lifetime history of psychotic disorder in self, report of psychotic disorder in first degree relative, history of alcohol-induced seizures, current mood disorder (past six months) or significant major medical disease, hypertension or diabetes, current pregnancy, mental retardation, left handedness, suicidal ideation, and positive urine drug screen.

Comprehensive substance abuse histories were obtained via a modified version of the Addiction Severity Index – Expanded (ASI-X; (McLellan et al., 1980)). All but four participants had been abstinent for at least the previous 30 days; all reported results remain significant if these four participants are removed from analyses. In addition, scores on the Beck Depression Inventory II (BDI-II; (Beck et al., 1996)), the Spielberger State-Trait Anxiety Scale (STAI; (Spielberger et al., 1983)), and the two-subtest Weschler Abbreviated Intelligence Scale for Adults (WASI; (Wechsler, 1999)) were collected and included as covariates in relevant analyses. Groups were carefully monitored for differences in gender, age, IQ and comorbid diagnoses, with targeted recruitment of individuals with specific characteristics sometimes employed to ensure no significant differences across important demographic and clinical variables (see Table 1 for full demographic/clinical details).

**Table 1.** Demographics and DSM-IV diagnoses across stimulant dependent individuals (SDI) and non-stimulant dependent individuals (non-SDI).

|  | **SDI** | **Control** | **Test** | **p** |
| --- | --- | --- | --- | --- |
|  |  |  | **Statistic** |  |
| n | 48 | 23 |  |  |
| Females | 10 (21%) | 1 (4%) | 2.09 | .15 |
| Age | 35.5 (8.3) | 28.9 (8.7) | 3.14 | .003 |
| Hispanic/Latino<sup>a</sup> | 26 (54%) | 11 (48%) | .06 | .81 |
| IQ | 102.8 (12.5) | 103.7 (11.8) | .30 | .77 |
| Mood Disorders |  |  |  |  |
| *Bipolar (Any)* | 0% | 0% |  |  |
| *Major Depression* | 13 (27%) | 3 (13%) | 1.04 | .31 |
| *Substance Induced Mood Disorder* | 5 (10%) | 0 (0%) | 1.23 | .27 |
| Schizophrenia/Psychotic |  |  |  |  |
| *Schizophrenia/Psychotic* | 0% | 0% |  |  |
| Substance-related |  |  |  |  |
| *Alcohol Dependence* | 29 (60%) | 7 (30%) | 4.57 | .03 |
| *Sedative/Hypnotic/Anxiolytic* | 1 (2%) | 0 (0%) | 0 | 1 |
| *Cannabis Dependence* | 23 (48%) | 9 (39%) | .19 | .66 |
| *Stimulant Dependence* | 23 (48%) | 0 (0%) | 14.18 | <.001 |
| *Opioid Dependence* | 21 (44%) | 5 (22%) | 2.37 | .12 |
| *Cocaine Dependence* | 40 (83%) | 0 (0%) | 40.57 | <.001 |
| *Hallucinogen/PCP* | 6 (12%) | 0 (0%) | 1.73 | .19 |
| *Poly Drug* | 3 (6%) | 0 (0%) | 0.35 | 1 |
| Anxiety |  |  |  |  |
| *Panic Disorder* | 7 (15%) | 0 (0%) | 2.26 | .13 |
| *Agoraphobia* | 1 (2%) | 0 (0%) | 0 | 1 |
| *Social Phobia* | 4 (8%) | 0 (0%) | .77 | .38 |
| *Specific Phobia* | 4 (8%) | 0 (0%) | .77 | .38 |
| *OCD* | 1 (2%) | 0 (0%) | 0 | 1 |
| *PTSD* | 14 (29%) | 2 (9%) | 2.65 | .10 |
| *Generalized Anxiety* | 5 (10%) | 0 (0%) | 1.23 | .27 |
| Beck Depression Inventory-II<sup>b</sup> | 15.39 (12.40) | 7.00 (7.27) | 2.94 | .006 |
| Cigarette Smoker | 19 (40%) | 8 (35%) | .15 | .70 |

The p value associated with the appropriate statistical test is included (t-test for age, z test of proportions for all other variables). <sup>a</sup>One control and two SDI failed to report ethnicity. <sup>b</sup>Two controls and seven SDI did not have BDI-II data available for analysis.

### Study Design

#### Time Estimation Task

The time estimation task was modified from (Shane and Weywadt, 2014; van der Veen et al., 2011), and designed to evaluate neural responses to the presentation of contingent performance feedback with and without variance associated with performance expectancies controlled. Each trial began with an asterisk, presented centrally on a video screen (visual angle ~3°). Participants were instructed to wait for the asterisk to disappear, and to press a button with their index finger when they felt one second had transpired from the asterisk’s offset. Next, participants rated their level of confidence in their estimate. Finally, following a jittered interval, participants received one of several feedback symbols regarding the accuracy of their estimate. Responses were deemed successful if they fell within a desired window surrounding 1000ms. This window was initially set at ± 250ms; thus time estimates between 750ms and 1250ms were initially deemed successful, and any estimate outside this window was deemed unsuccessful. The size of the window was adaptive, on a trial-by-trial basis, depending on participants’ performance throughout the course of the task. Every time a successful estimate was made, the window decreased by 30ms; every time an unsuccessful estimate was made, the window increased by 30ms. Previous work with this type of task has provided substantial evidence of the validity of this adaptive algorithm for balancing trial presentation in both healthy and clinical populations (Becker et al., 2014; Shane and Weywadt, 2014). The task was self-paced. Each participant completed 56 trials in each of two runs, for a total of 112 trials.

On half of the trials, feedback was *Informative*: participants received a ‘+’ sign to indicate an accurate estimate, or a ‘-’ sign to indicate an inaccurate estimate. On the other half of the trials, feedback was *Uninformative*: participants received a ‘?’ that provided no accuracy information. The ‘?’ feedback was presented randomly on 50% of accurate trials, and 50% of inaccurate trials; thus, on each trial, participants had an equal expectation of either *Informative* or *Uninformative* feedback. The uninformative feedback trials served as a critical design feature: unlike the *Incorrect_INFORMED_ > Correct_INFORMED_* contrast (within which performance and performance feedback are necessarily confounded), the *Incorrect_INFORMED_ > Incorrect_UNINFORMED_* contrast differs only in the type of feedback provided. Thus observed differences in neural response in the *Incorrect_INFORMED_ > Incorrect_UNINFORMED_* contrast can be specifically attributed to the presentation of the informed feedback.

#### MRI Acquisition

All participants completed a urine screen prior to scanning. MRI data was collected on a 3T Siemens Trio (Erlangen, Germany) whole body scanner with gradient-echo pulse sequence (TR=2000ms, TE=29, flip angle=75, 33 axial slices, 64 × 64 matrix, 3.75 × 3.75 mm2, 3.5 mm thickness, 1 mm gap) acquired with a 12-channel head coil.

### Data Analytic Strategies

#### Behavior Analyses

Behavioral measures were collected on each trial, and included estimation accuracy, estimation accuracy on trial n+1 (to evaluate post-feedback behavioral adjustments), and confidence ratings (on a 1-4 likert scale). Time estimations were extracted for each condition of interest. Time estimations greater than 10 seconds were removed, followed by all estimations greater than 3 standard deviations from individual subject means, to prevent undue influence from outliers. This process resulted in 104 total trials being removed from the analysis (1.4% of all data).

#### Neuroimaging Analyses

All image analyses were undertaken with FMRIB’s Software Library (FSL) version 4.1.0 (Smith et al., 2004), using standard preprocessing parameters. FSL’s Motion Correction using FMRIBs Linear Image Registration Tool (MCFLIRT; (Jenkinson et al., 2002)) was used to realign functional images within a given run to the middle volume within the run. Images were deskulled using FSL’s Brain Extraction Tool (BET; (Smith, 2002)), spatially smoothed with a 5 mm full-width half-max Gaussian kernel, temporally filtered using a high-pass filter of 50 sec, prewhitened using FMRIBs Improved Linear Model (FILM) and grand mean intensity normalized; all of these steps were performed using FMRIB’s Expert Analysis Tool (FEAT;(Woolrich et al., 2004)).

All analyses were run as part of a 3-stage process. In the first stage, customized regressors were created for each participant for four events: cue, estimate (correct, incorrect), confidence ratings, and feedback (informed-correct, informed-incorrect, uninformed-correct, uninformed-incorrect). In addition, six first-order motion parameters were included in the statistical model. Statistical analyses were performed using the general linear model as implemented in FEAT. Time series analyses were conducted using FILM (Woolrich et al., 2004) with local autocorrelation estimation. This first level analysis generated parameter estimates for each condition of interest. Contrast maps were registered to the participant’s high-resolution anatomical image and the MNI 152 brain template using FLIRT (Jenkinson et al., 2002). Next, analyses from each run within a participant were combined using a fixed effects model in FEAT. This second level analysis generated mean effects and within subject variance estimates that were used in third level analyses. Within the task, we were interested in 4 conditions of interest (i.e. Informed Correct, Informed Incorrect, Uninformed Correct, Uninformed Incorrect), which were analyzed using a 2 (Performance: Correct, Incorrect) × 2 (Feedback Type: Informed, Uninformed) × 2 (Group: SDI, non-SDI) mixed-model ANOVA. In addition, two primary contrasts were computed at the individual subject level to support hypothesis driven analyses of group differences in feedback processing: *Incorrect_INFORMED_ > Correct_INFORMED_* (within which performance expectancies necessarily varied), and *Incorrect_INFORMED_ > Incorrect_UNINFORMED_* (within which performance expectancies were effectively controlled); each of these contrasts were compared across groups using an independent samples t-test.

Whole brain group analyses were conducted using permutation testing with the randomise program within FSL. Threshold free cluster enhancement (TFCE) was used to correct for multiple comparisons at p < 0.05.

#### Years of use

Lifetime stimulant use assessment was consistent with general PhenX guidelines (Conway et al., 2014), and total years of use equaled the summed total years of use for all stimulants queried via the ASI-X. Correlation analyses investigated the relationship between lifetime years of use and percent signal change within each of two 10mm spherical ROIs drawn around the peak dACC voxels within the two clusters that emerged in the comparison of SDI and non-SDI for the *Incorrect_INFORMED_ > Incorrect_UNINFORMED_* contrast.

## Results

### Sample Characterization

Following exclusions for motion, performance and psychotropic drug use (see Supplement for full details regarding exclusion criteria), 71 participants (48 SDIs and 23 non-SDIs) were included in the final dataset. Table 1 provides full demographics.

### Behavioral Results

#### Accuracy

Necessarily, a main effect of Accuracy indicated that participants’ time estimates were more accurate on correct than incorrect trials, *F*(1, 69) = 414.38, *p* < 0.001. Importantly, however, the main effect of Information did not reach significance, indicating that performance was similar across *Informed* and *Uninformed* trials. Similarly, no group effects, nor interactions, reached significance (see Table 2 and Figure S1a in Supplement).

**Table 2.** Means and standard deviations in for performance measures in the time estimation task.

|  |  | SDI |  | Non-SDI |  |
| --- | --- | --- | --- | --- | --- |
| *Time estimation* |  | Informative | Uninformative | Informative | Uninformative |
| *Deviation from 1000 msec* |  |  |  |  |  |
|  | Correct | 143 (65) | 146 (78) | 158 (76) | 154 (83) |
|  | Incorrect | 432 (137) | 425 (136) | 425 (168) | 432 (202) |
| *Post-feedback Change* |  |  |  |  |  |
|  | Correct | 167 (64) | 182 (83) | 157 (77) | 160 (60) |
|  | Incorrect | 225 (81) | 210 (71) | 231 (116) | 209 (143) |

Deviation from 1000 milliseconds and post-feedback change values are presented in milliseconds.
Deviation scores represent the difference between the 1000 millisecond target estimation and the mean absolute value of deviations from one second. Post-feedback change scores represent the change in magnitude of deviation for trial $n$ to $n+1$ . To compute change, the difference in the absolute value of the deviation from 1 second on trial ( $n$ ) to the absolute value of the deviation from 1 second on trial ( $n + 1$ ) was computed. Main effects for performance (i.e. Correct vs. Incorrect) were significant, but main effects for Feedback Type (Informative vs. Uninformative) and group (SDI vs. non-SDI) were not significant.

#### Confidence

The time estimation task was specifically designed to maximize neural response to informative feedback by minimizing the development of outcome awareness. Consistent with this goal, while participants showed fairly high confidence in their estimation attempts (mean confidence rating = 1.78 (0.87)), the correlation between confidence and accuracy, while highly significant, was only low-to-moderate, *r*(68) = 0.14, *p* < .01. Importantly, this was true for both SDI (*r* = .17, *p* < .01) and non-SDI (*r* = .10, *p* < .01) groups (between group t-test: *t*(67) = 1.61, *p* = *ns*); moreover, no significant group differences in confidence were found in any condition (all *p*’s > .20).

#### Post-feedback Behavioral Adjustments

Trial-to-trial adjustments in estimation attempts were evaluated by calculating the deviation from 1000 ms on trial n+1. A 2 (Accuracy) × 2 (Information) × 2 (Group) mixed-model ANOVA indicated that participants’ estimation adjustments were greater following incorrect compared to correct trials, *F*(1, 69) = 62.94, *p* < 0.001. While main effects of Information and Group did not reach significance, a significant Accuracy × Information interaction, *F*(1, 69) = 3.86, *p* = 0.05, indicated that the difference between performance adjustments following correct/incorrect feedback was greater for *Informed* compared to *Uninformed* feedback (see Table 2 and Figure S1b in Supplement).

### Neuroimaging Results

#### Main Effects and Interactions

The 2 (Accuracy) × 2 (Information) × 2 (Group) mixed-model ANOVA revealed a main effect of Accuracy, within several clusters including dorsal and pregenual ACC d/pg ACC, bilateral nucleus accumbens, and bilateral occipital cortex. A main effect of Information was also identified within several regions including d/pg ACC, bilateral anterior insula, and nucleus accumbens. Finally, a main effect of group was identified within right cerebellum and left lingual gyrus. An Accuracy × Feedback interaction revealed significant clusters within bilateral occipital pole, ventral striatum, and left inferior parietal lobe. The Information × Group interaction, Group × Accuracy interaction and three-way Accuracy × Information × Group interaction revealed no significant clusters. See Table S1 in Supplement for all regions reaching significance within these higher-order analyses

#### Contrasts of interest

*Incorrect_INFORMED_ > Incorrect_UNINFORMED_* and *Incorrect_INFORMED_* > *Correct_INFORMED_* served as primary contrasts of interest. Here we report effects within these contrasts across all participants. Below we present the group-relevant distinctions. Other contrast-level effects can be found in the Supplement. *Incorrect_INFORMED_ > Incorrect_UNINFORMED_.* By controlling for expectancy effects related to differential performance, this contrast provides a well-controlled evaluation of the brain’s response to the specific presentation of negative feedback. Results revealed greater response following the presentation of *Incorrect_INFORMED_* feedback within several regions, including d/pg ACC and bilateral insula (see Figure 1). No regions showed greater activity following the presentation of *Incorrect_UNINFORMED_* feedback within this contrast.

**Figure 1.**
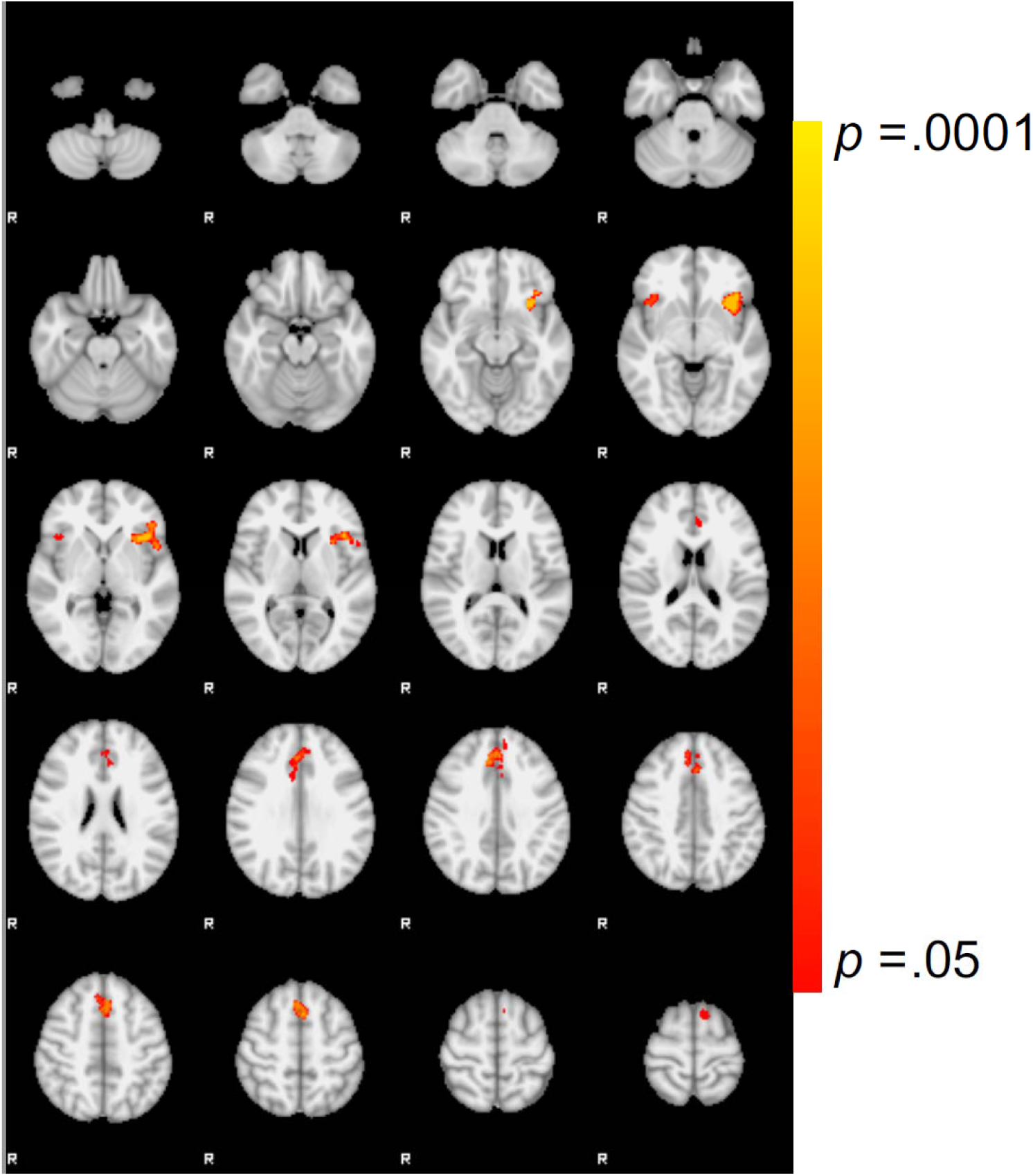
*Incorrect_INFORMED_ > Incorrect_UNINFORMED_* feedback. Main effect of *Incorrect_INFORMED_ > Incorrect_UNINFORMED_* (TFCE corrected at p < .05).

*Incorrect_INFORMED_* > *Correct_INFORMED_*. No regions showed greater response following *Incorrect_INFORMED_* feedback in this contrast. However, several regions, including bilateral insula, bilateral ventral striatum and pgACC, showed greater response to *Correct_INFORMED_* feedback (see Table 3).

**Table 3.** Brain regions showing significant differences in the primary contrasts of interest in the time estimation task. Images were corrected for multiple comparisons using TFCE with 5000 permutations at p < .05.

| Contrast | Region | Voxels | x | y | z | p |
| --- | --- | --- | --- | --- | --- | --- |
| *Incorrect*<sub>INFORMED</sub> > *Incorrect*<sub>UNINFORMED</sub> | preSMA/SFG | 851 | -2 | 14 | 52 | 0.018 |
|  | L OFC/Insula | 727 | -38 | 22 | -8 | 0.01 |
|  | R OFC/Insula | 124 | 44 | 20 | -6 | 0.033 |
| *Correct*<sub>INFORMED</sub> > *Incorrect*<sub>INFORMED</sub> | R OFC/Insula/IFG | 4528 | -32 | 18 | -14 | <0.001 |
|  | L OFC/Insula/IFG | 3112 | 34 | 20 | -16 | <0.001 |
|  | dACC/pgACC | 2771 | -4 | 36 | 24 | 0.001 |
|  | L IPL/Angular gyrus | 1181 | -52 | -56 | 40 | 0.001 |
|  | R IPL/Angular gyrus | 663 | 54 | -46 | 46 | 0.007 |
|  | L Occipital Pole | 383 | -24 | -98 | -8 | 0.002 |
|  | R Occipital Pole | 135 | 28 | -92 | -4 | 0.011 |
|  | R Middle Frontal/IFG | 7 | 62 | 22 | 26 | 0.049 |
| Non-SDI > SDI (*Incorrect*<sub>INFORMED</sub> > *Incorrect*<sub>UNINFORMED</sub>) |  |  |  |  |  |  |
|  | pgACC | 71 | 4 | 40 | 0 | 0.032 |
|  | Frontal pole | 54 | 42 | 56 | 4 | 0.037 |
|  | dACC | 46 | 6 | 36 | 20 | 0.043 |

#### Group Effects (SDI vs. non-SDI)

*A priori* hypotheses centered around differences in post-error neural responses between the SDI and non-SDI groups. To this end, we undertook targeted two-sample t-tests to evaluate group differences in the *Incorrect_INFORMED_ > Incorrect_UNINFORMED_* and *Incorrect_INFORMED_ > Incorrect_UNINFORMED_* contrasts. These analyses indicated that SDIs showed reduced ACC response compared to non-SDIs in the *Incorrect_INFORMED_ > Incorrect_UNINFORMED_* contrast in clusters within both dorsal and pregenual portions of the ACC (see Figure 2a, b; TFCE corrected p < 0.05). However, no such group differences were revealed in the *Incorrect_INFORMED_ > Correct_INFORMED_* contrast. Thus, only when performance expectancies were controlled did SDIs show attenuated ACC response compared to non-SDIs following presentation of negative feedback. Importantly, repeating this group level analysis with relevant covariates included (i.e. drug use, BDI, STAI, and age) did not alter the direction of these effects. (see *Supplement* for details).

**Figure 2.**
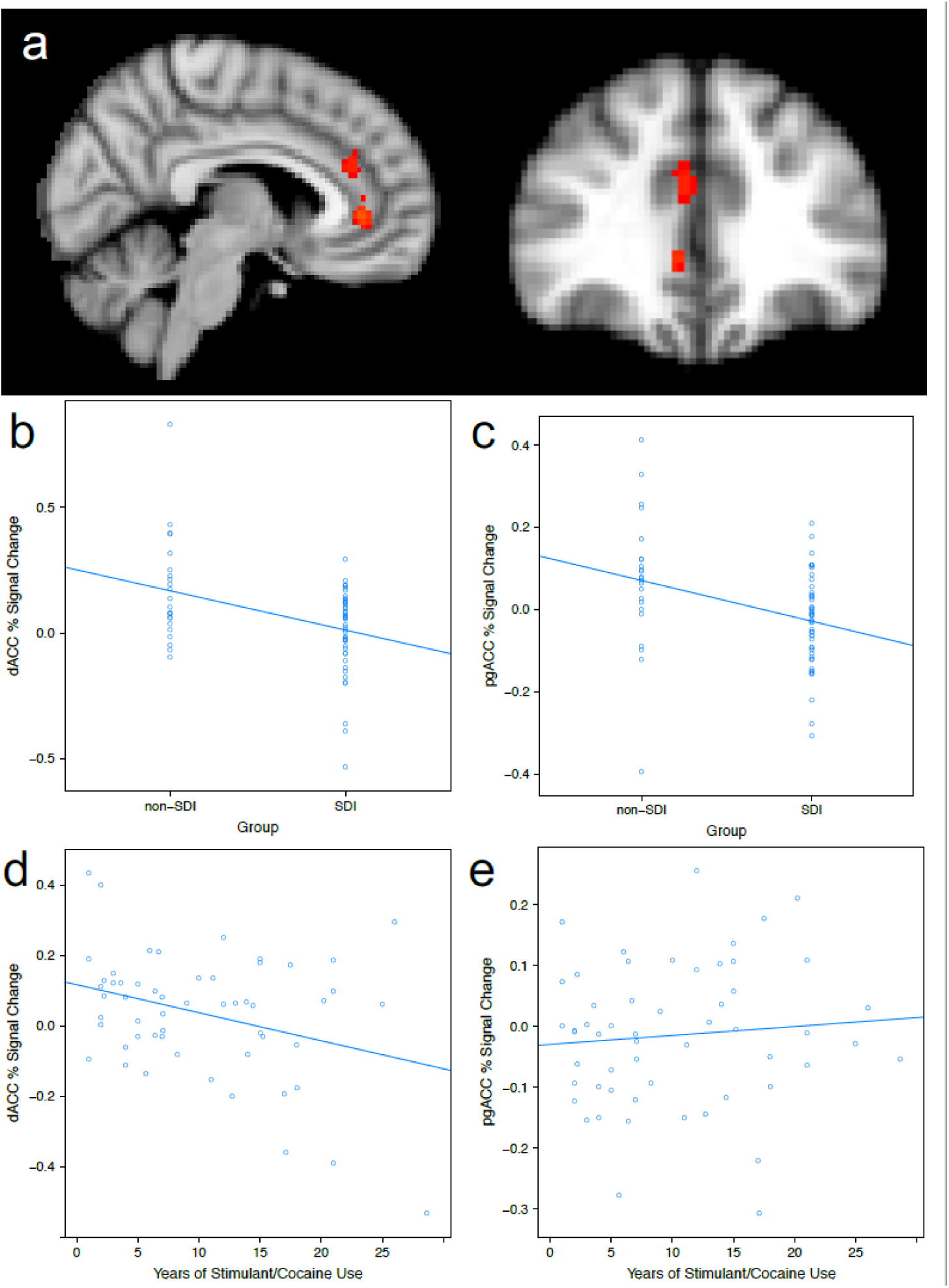
Association between error-related response, stimulant dependence and use history. a) dACC region within which controls (non-SDI; n=23) show greater response than stimulant dependent individuals (SDI; n=48) in the *Incorrect_INFORMED_ > Incorrect_UNINFORMED_* contrast (TFCE corrected at p <.05). Percent signal change in b) dACC and c) pgACC by group. Negative correlation between years of stimulant use and BOLD response during the *Incorrect_INFORMED_ > Incorrect_UNINFORMED_* contrast in the d) dACC and e) pgACC.

Group differences were also evaluated in the *Correct_INFORMED_ > Incorrect_INFORMED_* and the *Correct_INFORMED_ > Correct_UNINFORMED_* contrasts, to evaluate potential differences associated with reward processing. However, only one region within left lateral occipital cortex reached significance in the *Correct_INFORMED_ > Incorrect_INFORMED_* contrast (and none reached significance in the *Correct_INFORMED_ > Correct_UNINFORMED_* contrast. Thus, while broad dACC activation was seen to both positive and error feedback, only error-related responses manifested as group differences.

#### Relationship with Years of Use

Lifetime years of stimulant use was correlated with percent signal change in the dACC ROI derived from the group difference analysis. This analysis revealed a negative association between lifetime use and dACC response in the *Incorrect_INFORMED_ > Incorrect_UNINFORMED_* contrast, *r*(53) = -0.33, *p* < 0.02, (see Figure 2c). This correlation remained after controlling for IQ and lifetime years of other drug use, and dips slightly to *p* = .13 when controlling for age., Parallel analyses with the pregenual region of the ACC that emerged in the group analysis showed no such correlations with years of use.

## Discussion

The current study compared neural responses following the presentation of exogenous feedback in individuals with and without a stimulant dependence disorder. Results indicated that SDIs exhibited reduced dACC and pgACC response following the presentation of negative feedback; moreover, the magnitude of SDIs’ dACC reductions were correlated with the length of stimulant use history. While these effects were minimized after taking age into account, these findings converge with a growing body of work indicating that SDIs are characterized by identifiable error-related processing abnormalities (Franken et al., 2007; Hester et al., 2013; 2007 ; Kaufman et al., 2003; Patel et al., 2013) and emphasize the role that these abnormalities may play in the development and maintenance of stimulant use behaviors. Consistent with the proposed role of dACC in signaling to lateral prefrontal regions the need for phasic increases in cognitive control (Botvinick et al., 2001; Holroyd and Coles, 2002), a reduced dACC response to error feedback in SDIs may signify insufficient engagement of control mechanisms in the face of relevant error-related information.

The present study’s primary goal was to better characterize the extent to which SDIs’ error-processing abnormalities were the result of reduced awareness, or reduced sensitivity, to error-related information. While prior studies have sought similar goals, an inability to control for outcome expectancies has hindered a full understanding of the pathophysiology of SDIs’ error processing abnormalities. Importantly, the present results demonstrated that SDIs’ attenuated d/pg ACC response following negative feedback occurred only when variance associated with prior behavior (and thus behavioral expectancies) was fully controlled. This effect is subtle, but may provide clarification to a nascent, and currently inconsistent, literature (Parvaz et al., 2015; Patel et al., 2013). By controlling for expectancy effects, the task affords a pure measure of sensitivity to the mere presentation of negative feedback. In this context, SDIs showed a significantly attenuated dACC response. The fact that this effect disappeared when expectancies were allowed to vary further suggests that this reduced sensitivity may exist within the context of otherwise intact expectancy formation (see also (Parvaz et al., 2015)).

Of equal import, associations with SDIs’ lifetime stimulant use were also identified only when performance metrics were held constant. While this effect was mitigated after controlling for age, it nonetheless suggests that SDIs’ attenuated responses following presentation of negative feedback may in fact play an important role in the pathophysiology of the disease state. A handful of previous studies have reported similar associations between error-related responses and clinically-relevant abuse metrics (Luo et al., 2013; Marhe et al., 2013; Moeller et al., 2014; Steele et al., 2014); however, the driving force behind these associations has not yet been characterized. The present results suggest that a core insensitivity to the presentation of negative feedback may be an important factor in this process. One possibility is that a reduced sensitivity to negative feedback may interfere with the ability to incorporate the information held within that feedback into existing behavioral repertoires (Shane and Peterson, 2004).

It is relevant to note that both groups showed increased dACC response following presentation of positive feedback, but that these activations did not manifest as group differences. Thus, group differences in d/pgACC response were specific to the presentation of error-related feedback. While coinciding with a growing body of work that has positioned ACC as a core structure within the mesocorticolimbic reward system, it does conflict with some previous work that has reported reward-related abnormalities in SDIs and drug use behaviors (e.g. (Baker et al., 2016)). Much of this previous work has focused on evaluating cortical and sub-cortical responses to drug- and non-drug rewards (Goldstein et al., 2009; Volkow et al., 2011), and to engagement of cognitive control mechanisms towards inhibition of a desired reward (Garavan and Hester, 2007; Motzkin et al., 2014; Volkow et al., 2010). It may be that engagement of dACC in these inhibitory contexts are also abnormal in substance abusers; however, the mere presentation of positive feedback in the present study did not elicit similar group differences.

Some limitations of this study should be noted. First, despite our best attempts, the frequency with which SDI and non-SDI participants expressed comorbid dependencies differed somewhat across the two groups. While such differential comorbidities could have contributed to differences between SDI and non-SDI groups (Hester et al., 2007; Luo et al., 2013), it is important to note that all reported results remained significant when lifetime years of other drugs was included as a covariate (see supplementary results). Second, because both SDI and non-SDI groups were recruited through probation/parole, they all had significant criminal histories. To some extent, this may be expected, given the high rate of comorbidity between substance use disorders and criminal activity (Fazel et al., 2006). Nonetheless, it is important to note that the extent to which our results may generalize to SDIs without significant criminal histories, or less antisocial personalities (Cope et al., 2012) remains an open question. Third, we did not observe a similar relationship between dACC response and magnitude of post-feedback behavioral change. We note that while correlations between error-related responses and post-error changes are commonly reported within healthy populations (Danielmeier and Ullsperger, 2011), they are less reliably elicited within clinical populations including SDIs (Hester et al., 2007; Moeller et al., 2014). One possibility is that the effects of reduced error processing sum cumulatively, such that each indication of reduced processing may impose only minimal influence on subsequent behavior. Finally, as with most studies of this nature, it is difficult to rule out the possibility that participants’ dACC attenuations predated their stimulant use. Nonetheless, the fact that lifetime use correlated with magnitude of these attenuations suggests that longer-term, or more severe, use may further intensify alterations to this feedback processing system.

In summary, the current study demonstrated that dACC response following presentation of negative feedback, when unconfounded by error expectancies, was 1) attenuated in SDIs and 2) negatively associated with lifetime years of use. Negative feedback-related ACC response may provide a useful biomarker of stimulant dependence, and a potential target for treatments.

## Acknowledgements

This work was supported by a grant from the National Institute on Drug Abuse (R01DA026932) to Dr. Shane. The National Institute on Drug Abuse did not have any role in the design and conduct of the study; collection, management, analysis, and interpretation of the data; or preparation, review, or approval of the manuscript. We would like to acknowledge Mr. Ben Wasserott and Program Evaluation Services at the Center on Alcoholism, Substance Abuse, and Addictions at the University of New Mexico for recruitment of participants and data collection.

